# A newly-evolved chimeric lysin motif receptor-like kinase in *Medicago truncatula* spp. *tricycla* R108 extends its Rhizobia symbiotic partnership

**DOI:** 10.1101/2021.11.23.469708

**Authors:** Thi-Bich Luu, Anna Ourth, Cécile Pouzet, Nicolas Pauly, Julie Cullimore

## Abstract

- Rhizobial lipochitooligosaccharidic Nod factors (NFs), specified by *nod* genes, are the primary determinants of host specificity in the legume-Rhizobia symbiosis.
- We examined the nodulation ability of *Medicago truncatula* cv Jemalong A17 and *M. truncatula* ssp. *tricycla* R108 with the *Sinorhizobium meliloti nodF/nodL* mutant, which produces modified NFs. We then applied genetic and functional approaches to study the genetic basis and mechanism of nodulation of R108 by this mutant.
- We show that the *nodF/nodL* mutant can nodulate R108 but not A17. Using genomics and reverse genetics, we identified a newly-evolved, chimeric LysM receptor-like kinase gene in R108, *LYK2bis*, which is responsible for the phenotype and can allow A17 to gain nodulation with the *nodF/nodL* mutant. We found that LYK2bis is involved in nodulation by mutants producing non-*O*-acetylated NFs and interacts with the key receptor protein NFP. Many, but not all natural *S. meliloti* and *S. medicae* strains tested require *LYK2bis* for efficient nodulation of R108.
- Our findings reveal that a newly-evolved gene in R108, *LYK2bis*, extends nodulation specificity to mutants producing non-*O*-acetylated NFs and is important for nodulation by many natural *Sinorhizobia*. Evolution of this gene may present an adaptive advantage to allow nodulation by a greater variety of strains.

## Introduction

The importance of legumes in agriculture and the environment is largely due to their ability to form a N_2_-fixing symbiosis with Rhizobia bacteria, allowing them to grow independently of an additional nitrogen source (Graham & Vance, 2003). This ability has stimulated research on the evolution of such a successful mutualism, and the mechanisms that allow it to develop (Oldroyd *et al*., 2011; Griesmann *et al*., 2018). Legumes represent a huge family and show considerable variation in their specificity for their bacterial symbiont (Andrews & Andrews, 2017). Indeed, Rhizobia are polyphyletic, defined primarily by their ability to induce root nodules on their legume host, in which they differentiate into N_2_-fixing forms. Different genera/species/strains may show either a narrow or promiscuous host range (Masson-Boivin *et al*., 2009). Similarly, some legumes are highly promiscuous whereas others show exquisite fidelity to a narrow range of rhizobia (Andrews & Andrews, 2017; Chen *et al*., 2021).

Coevolution has ensured that Legume-Rhizobia couples are highly adapted to each other and has led to improvement of plant and bacterial fitness (Younginger & Friesen, 2019). However, as Rhizobia are horizontally transmitted symbionts, host-partner coevolution would not be possible if there were not signalling mechanisms to select the evolved partner from the external microbial milieu. Thus, partner choice signalling is a crucial step in the evolution of a successful symbiosis (Younginger & Friesen, 2019). Nod factors (NFs) are key rhizobial signals for initiating and maintaining the interaction between most well-studied legume-Rhizobia couples (Krönauer & Radutoiu, 2021). NFs are lipochitooligosaccharides (LCOs) with a basic structure of 3-5 *N*-acetyl glucosamine residues, *N*-acylated on the terminal non-reducing sugar. Various chemical substitutions on the basic structure lead to the primary mechanism of partner specificity (D’Haeze & Holsters, 2002; Oldroyd *et al*., 2011; Krönauer & Radutoiu, 2021). Although other signals may be important for some legume-Rhizobia interactions, such as cell-surface polysaccharides or protein effectors (Wang *et al*., 2018; Chen *et al*., 2021), the genomic linkage of the NF (*nod*) genes to nitrogen fixation (*nif*) genes (generally on a symbiotic megaplasmid or a genomic island), provides a means by which NF signalling can be used by the legume host to select its co-evolved and beneficial nitrogen-fixing partner (Younginger & Friesen, 2019).

On the plant side, lysin motif receptor like kinase (LysM-RLK) genes have evolved as key elements of NF perception and signalling. They encode plasma-membrane located proteins with three lysin motifs in the external region connected through a single transmembrane spanning domain to an internal kinase-like domain. These genes are involved in defence and mycorrhiza signalling in many plants, but the family is particularly expanded in legumes, where two genes show neofunctionalization to play essential roles in NF perception and nodulation (Buendia *et al*., 2018; Chiu & Paszkowski, 2020). One of these genes encodes a receptor with an inactive kinase domain and is known as *NFP/NFR5* (according to the species), whereas the other gene encodes a receptor with an active kinase domain and is known as *LYK3/NFR1* (Krönauer & Radutoiu, 2021). This latter gene evolved by tandem duplication, prior to the speciation of the legume family and it has been suggested that its neofunctionalization was fundamental to the evolutionary gain of a legume-specific NF receptor (De Mita *et al*., 2014). Further, more recent duplications of this and other genes in the cluster have led to a species-specific source of variation in NF signalling (De Mita *et al*., 2014; Sulima *et al*., 2017).

*Medicago truncatula* Gaertn. (closely related to alfalfa and pea), together with *Lotus japonicus*, is a widely used model for studying the molecular mechanisms leading to the Rhizobial symbiosis and also, more generally, for traits important for the successful use of legumes in agriculture (Garmier *et al*., 2017). Initially most work used the genotype *M. truncatula* ssp. *truncatula* cv Jemalong line A17 (referred to here as A17), but more recently, genomic resources have been developed for many other genotypes (see the Hapmap project – https://medicagohapmap2.org/) in order to exploit the considerable genetic variation of this species for these studies (Garmier *et al*., 2017). Of note, *M. truncatula* ssp. *tricycla* line R108-1 (referred to as R108) has achieved considerable success due to the development of a *Tnt1*-insertion population resource and its superior genetic transformation properties (Garmier *et al*., 2017; Kaur *et al*., 2021). Studies on A17 have shown that it has a narrow partner specificity, being nodulated primarily by the *Sinorhizobium (*also known as *Ensifer)* species *S. meliloti* and *S. medicae* (Andrews & Andrews, 2017; Kazmierczak *et al*., 2017). Analysis of the widely-used *S. meliloti* strain 2011 has shown that it produces NFs that are 6-*O*-sulphated on the reducing sugar and contain predominantly a C16:2 acyl chain on the terminal non-reducing sugar, which may also be 6-*O*-acetylated. The presence of the sulphate (specified by the *nodH* gene) is essential for nodulation (Roche *et al*., 1991) and the modifications on the terminal non-reducing sugar are also important. For example, a double *nodF/nodL* mutant which couples mutations in *nodF* (leading to *N*-acylation mainly with a C18:1 fatty acid) and *nodL* (leading to lack of 6-*O*-acetylation), is almost completely Nod^-^, whereas a *nodF* or a *nodL* mutant induce similar or a reduced number of nodules (Ardourel *et al*., 1994; Smit *et al*., 2007). On the plant side, *NFP* is required for all NF responses and its role can be partially substituted by *NFP* genes from other species (Bensmihen *et al*., 2011; Girardin *et al*., 2019). Studies on a weak *lyk3* mutant of A17 (Smit *et al*., 2007) and recent structural studies (Bozsoki *et al*., 2020) suggest that *LYK3* is involved in the perception of specific NF features.

Comparative studies between A17 and R108 have shown that they are highly divergent members of the *M. truncatula* species (Zhou *et al*., 2017), show differences in their symbiotic effectiveness with different Rhizobial strains (Kazmierczak *et al*., 2017) and exhibit differences in partner choice when confronted with a mixed *Sinorhizobium* (*Ensifer*) inoculum (Burghardt *et al*., 2018). In this article, we report that R108 has an extended NF-related strain specificity compared to A17 and have identified a newly evolved *LysM-RLK* gene, unique to R108, which is required for this phenotype.

## Materials and methods

### Seed germination and growth conditions

*Medicago* seeds were scarified in 95% sulphuric acid for 5 min, washed with distilled water two times and surface-sterilized in bleach (3.2% chlorine) for 3 min, then washed three times. After that, the seeds were soaked in water for 1 h and placed on 1% agar, supplemented with 1 μg ml^-1^ GA3. The plates were kept upside-down at 4°C for 5 d and then put at 16°C overnight for germination. Seedlings were transferred to pots and grown in a growth chamber at 22°C with a 16 h photoperiod.

### cDNA cloning

R108 roots were collected and immediately frozen in liquid N_2_. Total RNA was extracted using a Nucleospin plant RNA extraction kit (Marcherey-Nagel GmbH & Co. KG, Germany). cDNA was synthesized using SuperScript IV Reverse Transcriptase kit (Thermo Fisher Scientific, USA).

*LYK2bis* coding sequence (CDS) was amplified with Phusion High Fidelity DNA Polymerase (F530S, Thermo Fisher Scientific, USA) using R108 root cDNA as template and the following primers (5’-ATGAAACTAAAAAATGGCTTAC-3’ and 5’-TCTAGTTGACAACAAATTTATG-3’). The PCR product was cloned into pJet1.2/blunt using CloneJET PCR Cloning Kit (K1231, Thermo Fisher Scientific, USA). The correct clone was selected by sequencing, then used for further cloning steps.

### Constructs for in planta protein expression

For *M. truncatula* root transformation, full length CDS of *LYK2bis* and of *LYK3* from A17 (LYK3-A17) were fused at the C-terminus with 3xFLAG under the control of the Ubiquitin promoter from *L. japonicus* (ProLjUbi) by Golden gate cloning, using a vector based on pCAMBIA2200 expressing DsRED (Fliegmann *et al*., 2016). For *Nicotiana benthamiana* leaf agro-infiltration, full length CDS of LYK2bis and NFP were fused at the C-terminus with mCherry and GFP tag proteins, respectively, under the control of ProLjUbi. The vector pCambia2200ΔDsRED was used (Fliegmann *et al*., 2016).

### Mutants and nodulation tests

*Tnt1* insertional mutants, *lyk2-1R* (NF13076), *lyk2bis-1R* (NF15454) and *lyk3-1R* (NF2752) of R108, obtained from the Noble Research Institute (USA), were used for nodulation tests. R108 and A17 were used as wild-type (WT) controls. Note that “*R*” designates a R108 mutant.

Germinated seedlings were transferred into pots (8×8×8 cm^3^ in size), filled with sterilized attapulgite clay granules (Oil Dri, UK). To each pot, 80 ml of Fahraeus medium supplemented with 0.2 mM NH_4_NO_3,_ was added. After 3 d, plants were inoculated with approximately 10,000 bacteria/plant of *S. meliloti* 2011 WT or mutant strains (Table **S1**). The number of nodules per plant was counted at 21 dpi. When appropriate, bacterial infection was determined by LacZ straining with X-gal. All results were confirmed in separate biological experiments.

For the nodulation test with natural strains, germinated seedlings were transferred into test tubes with 20 ml agar slants of Fahraeus medium with 13 g/l of agar Kalys HP 696 (Kalys SA, Bermin, France), supplemented with 0.2 mM NH_4_NO_3_. After 5 d, plants were inoculated with approximately 10,000 bacteria/plant of different rhizobial strain. The lower part of the tubes was covered by brown paper to avoid excessive light access. The number of nodules per plant was analysed at 28 dpi. Results presented are from two separate biological replicates.

### Acetylene reduction assays

Assays were performed on inoculated plants at 28 dpi as described by Hardy *et al*., (1968). Briefly, 1 ml of acetylene was injected into a test tube containing one single inoculated plant and closed with a septum. Tubes were incubated in a growth chamber for two hours. Then, 400 μl of gas samples were analysed on an Agilent 7020 gas chromatograph, equipped with a flame ionization detector. Activity was normalised with the number of nodules per plant.

### Complementation and gain-of-function assays

Using *Agrobacterium rhizogenes*-mediated transformation (Boisson-Dernier *et al*., 2001), seedlings of *lyk2bis-1R* and A17 were transformed using strains containing either empty vector (EV) or ProLjUb:LYK2bis-3xFLAG or ProLjUb:LYK3-A17-3xFLAG constructs, and transformants were selected on medium containing 25 μg ml^-1^ kanamycin, and after two-weeks growth, by expression of the DsRed marker. R108 seedlings transformed with EV were used as control. Nodulation was analysed in pots as above after 4 wk. The number of nodulated plants and the number of nodules/nodulated plant were analysed.

### Agro-infiltration and Fluorescence Lifetime Imaging on Nicotiana benthamiana leaves

*Agrobacterium tumefaciens LBA4404* strains containing ProLjUb:LYK2bis-mCherry or NFP-GFP fusion constructs were used to agro-infiltrate the three to four oldest leaves of each *N. benthamiana* plant in which the OD_600_ of LYK2bis-mCherry was five times higher than that of NFP-GFP in order to have similar expression levels.

Two days post infiltration, the protein expression in leaves was assessed by confocal microscopy. Forster resonance energy transfer (FRET) between the fluorophores was analysed by Fluorescence Lifetime Imaging Measurements (FLIM) on a Leica TCS SP8 SMD which consists of an inverted LEICA DMi8 microscope equipped with a TCSPC system from PicoQuant. The excitation of the FITC donor at 470 nm was carried out by a picosecond pulsed diode laser at a repetition rate of 40 MHz, through an oil immersion objective (63×, N.A. 1.4). The emitted light was detected by a Leica HyD detector in the 500-550 nm emission range. Images were acquired with acquisition photons of up to 1500 per pixel.

From the fluorescence lifetime images, the decay curves were calculated per pixel and fitted (by Poissonian maximum likelihood estimation) with either a mono-or double-exponential decay model using the SymphoTime 64 software (PicoQuant, Germany). The mono-exponential model function was applied for donor samples with only GFP present. The double-exponential model function was used for samples containing GFP and mCherry. Experiments were repeated at least three times. The efficiency of energy transfer (*E*) based on the fluorescence lifetime (*τ*) was calculated as *E* = 1 − (*τ*_D+A_/*τ*_D−A_), where *τ*_D+A_ is donor fluorescence lifetime in the presence of acceptor while *τ*_D−A_ is the donor fluorescence lifetime in the absence of acceptor.

### Kinase assays

Glutathione-S-transferase (GST) tagged proteins of the predicted intracellular region of LYK2bis (termed the KD) were expressed in *E. coli* DH5a and the proteins purified using glutathione resin (GE Healthcare, USA) as described (Fliegmann *et al*., 2016). LYK2bis-KD was released from the resin using PreScission Protease (GE27-0843-01, Sigma Aldrich, Germany). LYK2bis-KD was incubated with kinase buffer containing ^32^P-ATP alone or with purified GST/NFP-KD, GST/LYR2-KD, GST/LYR3-KD, GST/LYR4-KD, GST/LYK3-deadKD (G334E mutation), Myelin Basic Protein (MyBP) or GST at 25°C for 1 h and the proteins analysed by SDS-PAGE, followed by Coomassie staining and Phosphor Imaging.

### Mycorrhization tests

Mycorrhization tests were performed using the gridline intersect method as described in Gibelin-Viala *et al*. (2019). Briefly, *lyk2bis-1R* and R108 seedlings were inoculated with 200 spores per plant of *Rhizophagus irregularis* DAOM197198 (Agronutrition, Toulouse, France) and colonisation was assessed at 3- and 5-wk post inoculation (wpi) using 15 plants/genotype/time-point.

### Bioinformatic analysis

The 1^st^ exon encoding the whole LysM domain of *LYK2* (640 bases), *LYK3* (637 bases) and *LYK2bis* (640 bases) from R108 were used with blastn to screen all *M. truncatula* genomes available in the Medicago BLAST Service of the Hapmap2 database (https://medicagohapmap2.org/). The first hit for each query was extracted. The percentages of identity (%ID) between hits found with the *LYK3-R108* probe and the *LYK2bis* sequence were calculated by EMBOSS Needle.

## Results

### R108 shows an extended nodulation specificity with a S. meliloti nodF/nodL mutant strain

To determine the nodulation specificity of the *M. truncatula* R108 genotype, seedlings of R108 and A17 were inoculated with either *S. meliloti* 2011 WT or its *nodF/nodL* mutant. The number of nodules was counted at 21 dpi. No nodules were found on A17 plants inoculated with *nodF/nodL*, which is consistent with the results on *M. truncatula* cv. Jemalong obtained previously (Ardourel *et al*., 1994) (Fig. **1**). However, R108 showed a similar nodulation capacity with both WT and *nodF/nodL* strains (Fig. **1**). Most (> 95%) of the R108 nodules with both strains were pink (suggesting expression of leghaemoglobin) and LacZ staining showed that they were well-infected with bacteria. Also, acetylene reduction assays suggest that the nodules from the two strains exhibit a similar ability to fix N_2_ (Fig. **S1**). These results suggest that R108 has an extended nodulation specificity, compared to A17, which includes the *nodF/nodL* mutant.

**Fig. 1.**
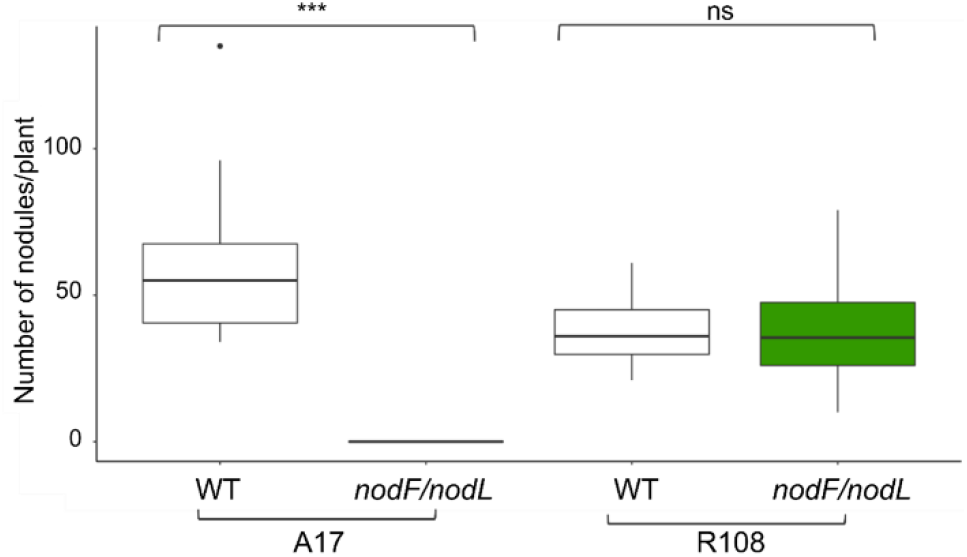
*Medicago truncatula* R108 is able to nodulate with the *Sinorhizobium meliloti nodF/nodL* strain. The number of nodules in 20 plants/genotype/inoculation was analysed at 21dpi. Statistical analyses were performed using Student’s *t*-test (ns, not significant; ***, *P* < 0.001). White shading is for plants inoculated with the WT strain; green shading is for plants inoculated with the *nodF/nodL* strain.

### A newly-evolved gene in R108, LYK2bis, is required for nodulation with the nodF/nodL strain

The perception of NF and nodulation in *M. truncatula* have been shown to require two LysM-RLKs, which are NFP and LYK3 in A17 (Arrighi *et al*., 2006; Smit *et al*., 2007). In the R108 genotype, NFP is reported to play essential roles for these processes (Feng *et al*., 2019), therefore, to understand the extension of nodulation in this ecotype, we focused on the *LYK* gene cluster, close to *LYK3*.

Using the genome sequence of R108 v1.0 (Kaur *et al*., 2021), we located the *LYK* gene cluster on chromosome 5, using the sequences of *LYK2* (MtrunA17Chr5g0439701) and *LYK3* (MtrunA17Chr5g0439631) from A17. Between *LYK2* and *LYK3*, an additional *LysM-RLK* gene was found, which is not present in A17, and is designated as *LYK2bis* (Fig. **2a**). Notably, the exon/intron organization is conserved among these genes with 12 exons in each (Fig. **2a**). The predicted amino acid sequence alignment between LYK2, LYK2bis and LYK3 proteins of A17 and R108 indicates that LYK2bis is a chimera with a similar extracellular domain to LYK2 up to the middle of LysM3 and then shares a highly conserved sequence with LYK3 (Fig. **2b**, Fig. **S2** and Table **S2**). In order to examine the presence of *LYK2bis* in *M. truncatula* ecotypes, the nucleotide sequence of the first exon of *LYK2, LYK3* and *LYK2bis* (containing the whole LysM region) were used to screen the 23 other *M. truncatula* genomes available for BLAST screening in the Hapmap project (https://medicagohapmap2.org/). Separate sequences corresponding to *LYK2* and *LYK3* were identified in all cases (Table **S3**). However, in genotype HM026, the available *LYK3* sequence seems to be not complete. The *LYK2bis* probe identified primarily the *LYK2* gene in all the genomes, with a lesser homology of this exon to the *LYK3* sequence. We did not find any evidence for an additional *LYK2* or *LYK3*-like gene in these genomes. These results suggest that *LYK2bis* is unique to R108 in these *M. truncatula* Hapmap genomes.

**Fig. 2.**
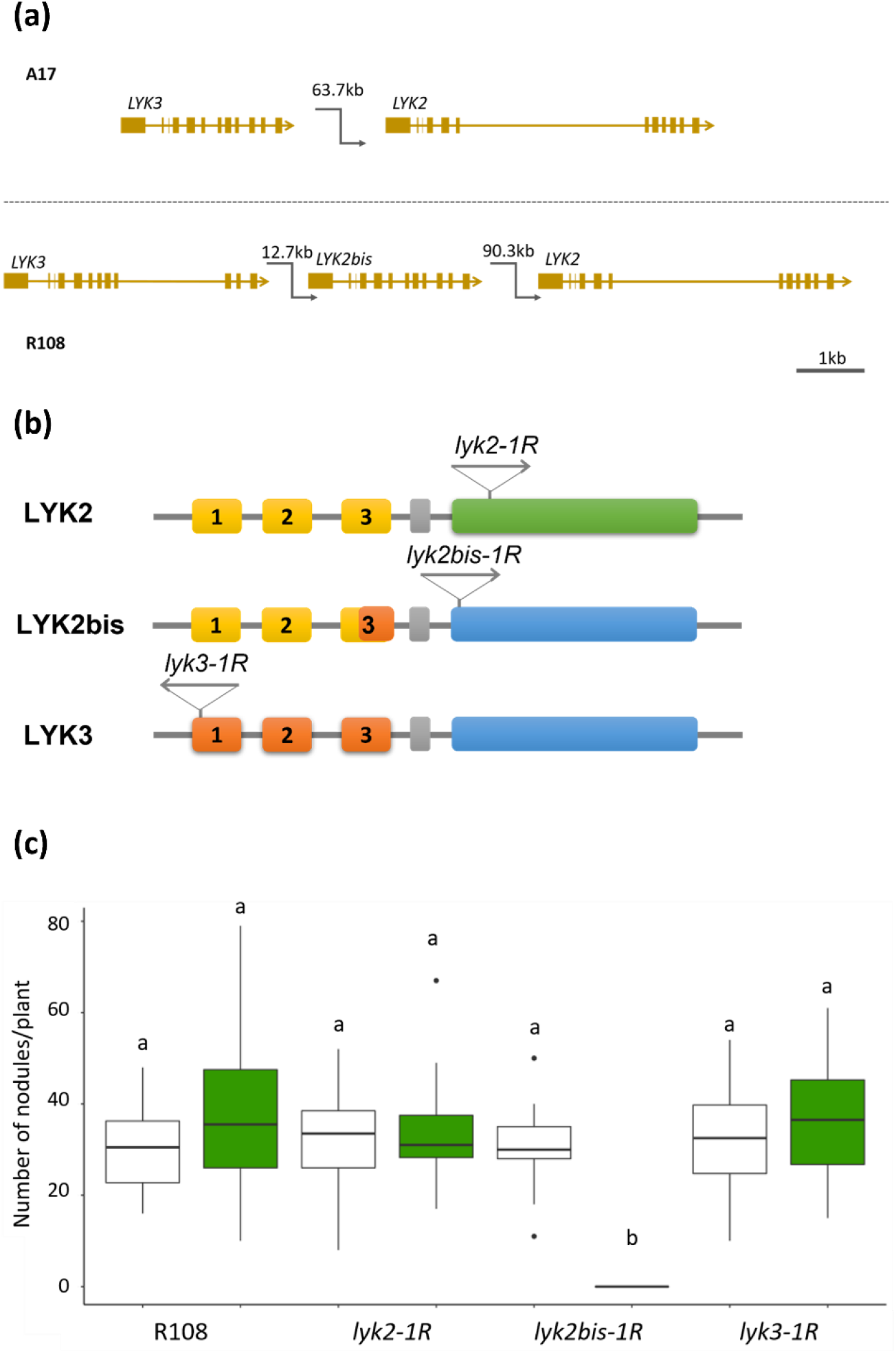
*LYK2bis* is a newly-evolved gene in *Medicago truncatula* R108 and is responsible for nodulation with the *Sinorhizobium meliloti nodF/nodL* strain. (a) Schematic representation of the *LYK3-LYK2* region in *M. truncatula* A17 and R108. Coding sequences are shown in filled blocks; Non-coding sequences are shown in lines. kb = kilobases. (b) Schematic representation of LYK2, LYK2bis and LYK3 proteins of R108, showing the structure of the proteins and the *Tnt1* insertional sites in *lyk2-1R, lyk2bis-1R* and *lyk3-1R mutants*. The proteins are predicted to have three extracellular LysM domains (yellow/orange), a transmembrane domain (grey) and an intracellular kinase domain (blue/green). (c) *LYK2bis* is responsible for nodulation with the *nodF/nodL* mutant in R108. *lyk2-1R, lyk2bis-1R* and *lyk3-1R* mutants were phenotyped for nodulation with either *S. meliloti* WT or the *nodF/nodL* mutant strain. 20 plants/genotype/inoculation were analysed at 21dpi. Statistical analyses were performed using One-way ANOVA (*P* < 0.05). Lowercase letters indicate significant difference. White shading is for plants inoculated with WT; Green shading is for plants inoculated with its *nodF/nodL* mutant.

To determine the roles of these genes in nodulation, *Tnt1* insertional mutants of *LYK2, LYK2bis* and *LYK3* in the R108 background were identified from the Noble Foundation resources (Fig. **2b**). The mutants were inoculated with either *S. meliloti* WT or *nodF/nodL* strains and phenotyped at 21 dpi. All tested lines formed a similar number of nodules with the WT strain (Fig. **2c**) indicating that, unlike the crucial role of *LYK3* in A17, none of those genes plays an essential role in nodulation of R108 with the *S. meliloti* WT strain. Similarly, there was no significant difference in the number of nodules formed in *lyk2-1R* and *lyk3-1R* mutants with the *nodF/nodL* strain, compared to R108. In contrast, no nodules formed on *lyk2bis-1R* plants inoculated with *nodF/nodL* suggesting that the *LYK2bis* gene is responsible for nodulation of R108 with the *nodF/nodL* mutant strain.

To validate the involvement of *LYK2bis* in nodulation with the *nodF/nodL* strain, we performed a complementation experiment of the *lyk2bis-1R* mutant. Using *A. rhizogenes*-mediated transformation, constructs of *LYK2bis, LYK3-A17* and empty vector (EV) were expressed in the *lyk2bis-1R* mutant. As shown in Fig. **3**, while neither *LYK3* nor EV transformed plants formed nodules with the *nodF/nodL* strain, nodulation was restored in most plants transformed with *LYK2bis*. This result confirms the essential role of *LYK2bis* in nodulation of R108 with the *nodF/nodL* mutant.

**Fig. 3.**
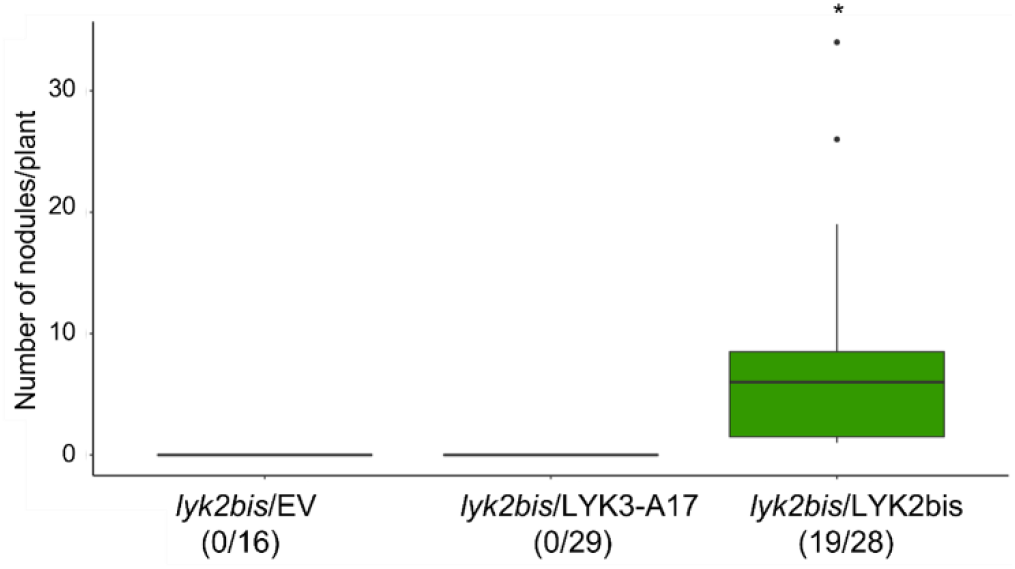
Complementation analysis confirms the role of *LYK2bis* in nodulation of *Medicago truncatula* R108 with the *Sinorhizobium meliloti nodF/nodL* strain. *lyk2bis-1R* roots were transformed with constructs of empty vector (EV), LYK3-A17 and LYK2bis using *A. rhizogenes*. Transformed plants were analyzed at 4 wpi. Statistical analyses were performed using Student’s *t*-test (*, *P* < 0.05). Numbers below indicate number of nodulated plants/total transformed plants.

### LYK2bis allows A17 to gain the ability to nodulate with the nodF/nodL strain

As A17 is not able to nodulate with the *nodF/nodL* strain and does not contain a *LYK2bis* gene in the genome, we performed a gain-of-function assay in A17 to determine whether introduction of *LYK2bis* is sufficient to gain nodulation with the *nodF/nodL* strain in this genotype. The experiment was done using a similar approach to the *lyk2bis-1R* complementation. Most of the A17 plants (29/36) transformed with the *LYK2bis* construct nodulated with *nodF/nodL* while none of the ones transformed with *LYK3-A17* and EV formed nodules (Fig. **4**). In a similar experiment but using the A17 *lyk3-1* mutant, 11/18 transformed plants formed nodules with the *nodF/nodL* mutant (data not shown). This evidence reveals that *LYK2bis* is central for interacting and nodulating with the *nodF/nodL* strain and can transfer this ability to another genotype, independently of *LYK3*.

**Fig. 4.**
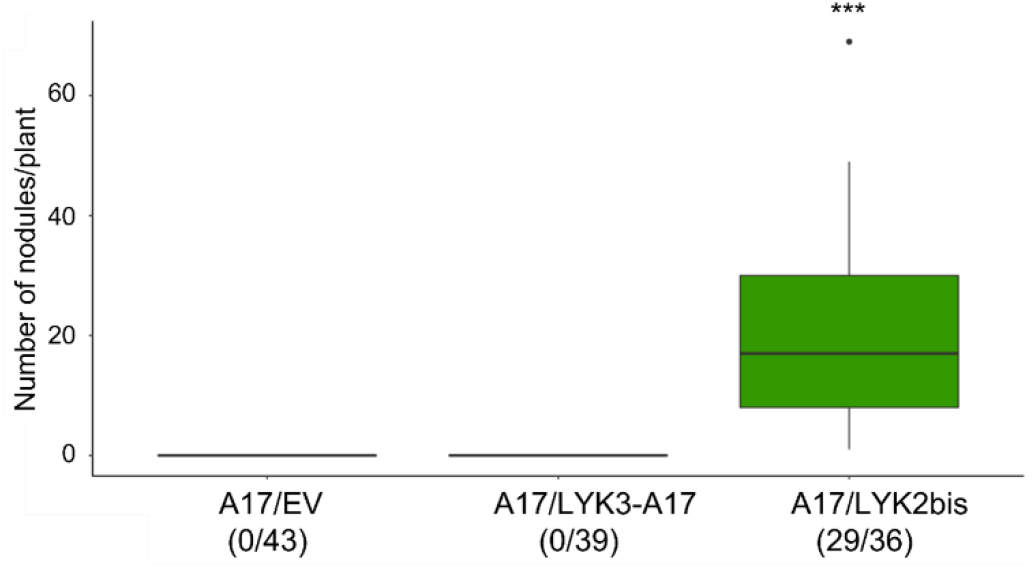
*LYK2bis* allows *Medicago truncatula* A17 to gain the ability to nodulate with the *Sinorhizobium meliloti nodF/nodL* strain. A17 roots were transformed with constructs of EV, LYK3-A17 and LYK2bis using *A. rhizogenes*. Transformed plants were analysed at 4 wpi. Statistical analyses were performed using Student’s *t*-test (***, *P*-value < 0.001). Numbers below indicate number of nodulated plants/total transformed plants.

### LYK2bis is involved in the perception of specific NF decorations leading to successful nodulation in R108

*S. meliloti nodF/nodL* produces NFs that differ from those of the WT strain in lacking the *O*-acetylation and containing C18:1 fatty acid chains rather than C16 on the terminal non-reducing sugar (Ardourel *et al*., 1994). The *nodL* mutant produces non-*O*-acetylated NFs (Ardourel *et al*., 1995), whereas mutants in *nodF* and the neighbouring *nodE* lead to NFs with the C18:1 acyl chains (Ardourel *et al*., 1994). We found that both these strains could form nodules on R108 as expected (Fig. **5**). However, while a similar number of nodules formed on *lyk2bis-1R* and R108 with the *nodFE* strain, no nodules were found on *lyk2bis-1R* inoculated with the *nodL* mutant (Fig. **5**).

**Fig. 5.**
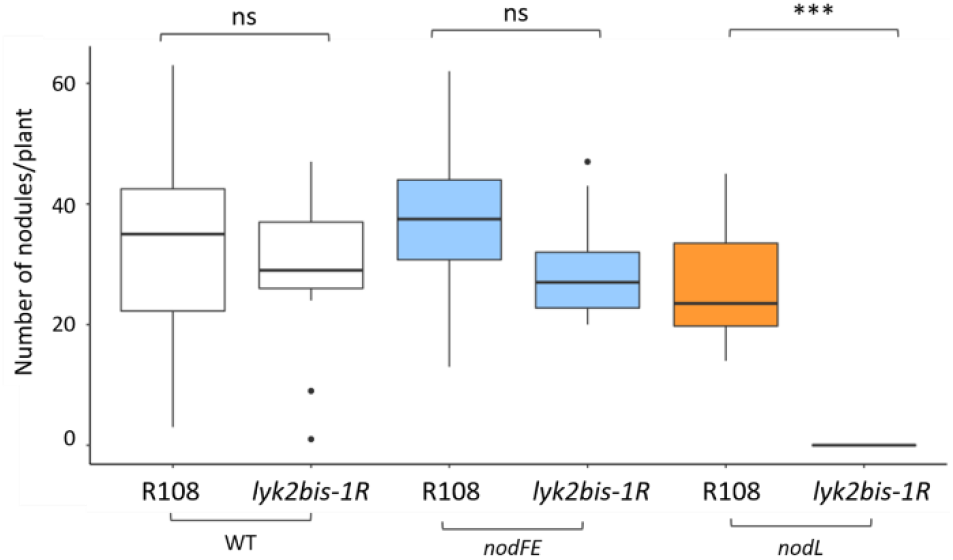
Nodulation specificity of the *Medicago truncatula lyk2bis-1R* mutant with different *Sinorhizobium meliloti* mutant strains. The *lyk2bis-1R* mutant was tested for nodulation with *nodF/nodE* and *nodL* mutant strains. The number of nodules were counted at 21 dpi. Statistical analyses were performed using Student’s *t*-test (ns, not significant; ***, *P* < 0.001). White shading is for plants inoculated with WT; Light blue shading is for plants inoculated with *nodF/nodE;* Orange shading is for plants inoculated with *nodL*.

Our results suggest that while other protein(s) can recognize rhizobia producing C18:1 NFs in R108, LYK2bis is particularly involved in the perception of rhizobia producing non-*O*-acetylated NFs.

### LYK2bis allows R108 to successfully nodulate with many natural rhizobial strains

To examine the ecological role of the newly-evolved gene *LYK2bis*, we performed nodulation tests on R108 and the *lyk2bis-1R* mutant with twenty-two natural *Sinorhizobium* strains (Table **S4** and Fig. **6**). The strains were selected as being representative of different multi-locus sequence types, or isolated from different *Medicago* species (Sugawara *et al*., 2013), including ones trapped from a soil in the South of France (Bailly *et al*., 2006). They also include strains from different geographical origins, particularly around the Mediterranean basin and including the Middle East. A preliminary experiment in tubes using a small number of plants showed that 9/14 of the *S. meliloti* strains and 5/8 of the *S. medicae* strains produced less nodules on the *lyk2bis* mutant than on R108 (*P* < 0.05). No correlation was seen between geographical origin and dependence on *LYK2bis*. To confirm the results, two replicate experiments were set up using a subset of 8 of these strains. All of the strains showed good nodulation of R108 but with some variability (from 8 to 20 nodules per plant in the tube system). Four of the strains however could barely nodulate the *lyk2bis* mutant (less than 1 nodule per plant) whereas 3 strains, like Sm 2011, showed no significant difference between the *lyk2bis* mutant and R108, in terms of nodule number. It is notable that strains of *S. meliloti* and *S. medicae* were represented in both the *LYK2bis*-dependent and - independent classes. These results show that *LYK2bis* is very important for nodulation by many but not all *S. meliloti* and *S. medicae* strains tested.

**Fig. 6.**
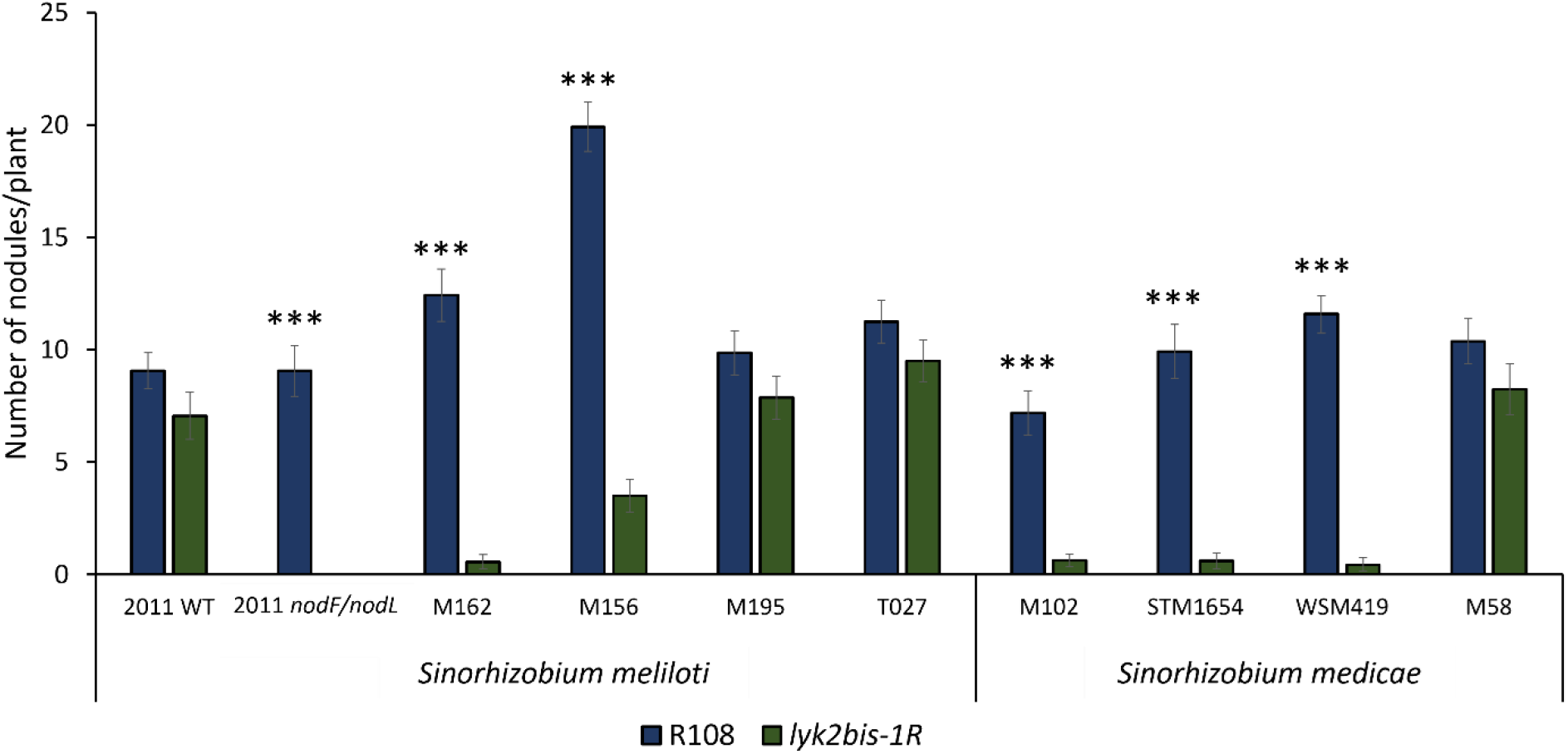
*LYK2bis* is required for R108 to successfully nodulate with some but not all natural *Sinorhizobium* strains. Seedlings of R108 and *lyk2bis-1R* mutant were inoculated with different strains in a tube system. 16 plants/strain for each genotype were analysed at 28 dpi. Statistical analyses between each R108/*lyk2bis* pair for each strain were performed using Student’s *t*-test (***, *P* < 0.001). Dark blue shading is for the R108 genotype and dark green shading is for *lyk2bis-1R*.

### LYK2bis does not play an important role in mycorrhization

As some Myc-factors have an identical structure to the major *nodF/nodL* NFs (Maillet *et al*., 2011), we also tested the role of *LYK2bis* in forming arbuscular mycorrhiza (AM). Seedlings of *lyk2bis-1R* and R108 were inoculated with spores of *Rhizophagus irregularis* and phenotyped for AM colonization at 3 wpi and 5 wpi. Both genotypes showed good mycorrhization at both time points (Fig. **7**), indicating that *LYK2bis* is not essential for mycorrhization.

**Fig. 7.**
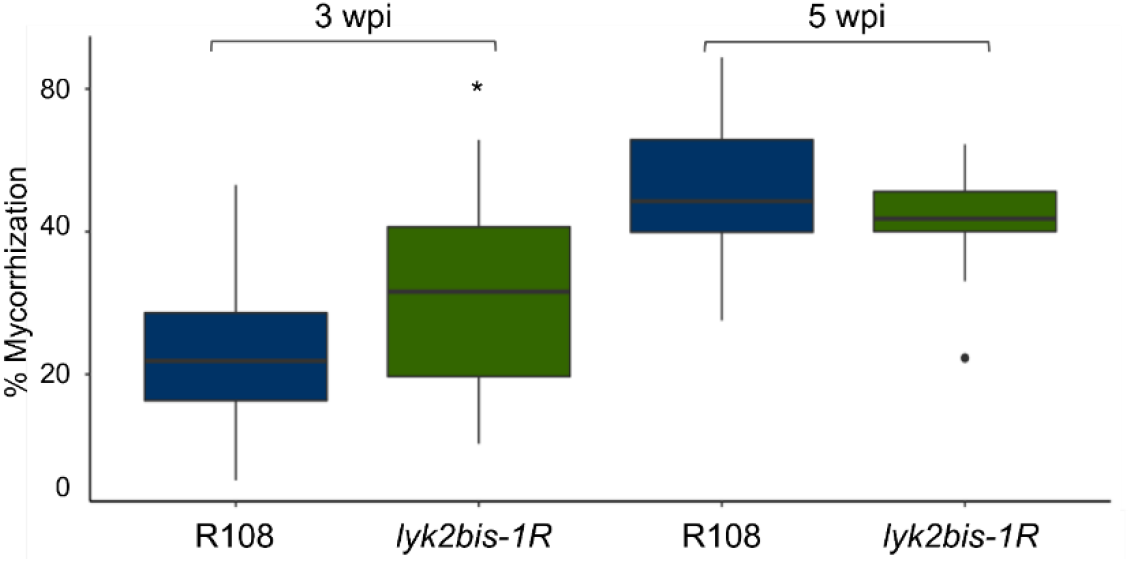
*LYK2bis* does not play an essential role in mycorrhization. Seedlings were inoculated with *Rhizophagus irregularis* and examined at 3 and 5 wpi. 15 plants/genotype at each time point were analysed. Statistical analyses were done using Students’ *t*-test (*, *P* < 0.05). Dark blue shading is for the R108 genotype and dark green shading is for *lyk2bis-1R*.

### LYK2bis physically interacts with NFP

NFP is required for nodulation in R108 (Feng *et al*., 2019) as well as in A17 (Arrighi *et al*., 2006), therefore, to test whether LYK2bis could be part of an NFP receptor complex, we investigated whether the two proteins interact using FRET-FLIM technology.

A construct of the LYK2bis protein fused with mCherry was co-expressed with a construct of a NFP-GFP fusion in *N. benthamiana* leaves using *A. tumefaciens*-mediated expression. Interestingly, at 3 dpi, all leaves expressing both proteins showed strong cell-death (Fig. **8a**) similar to that observed in the co-expression of NFP and LYK3-A17 (Pietraszewska-Bogiel *et al*., 2013). This suggests that LYK2bis may functionally interact with NFP in a similar mechanism to LYK3.

**Fig. 8.**
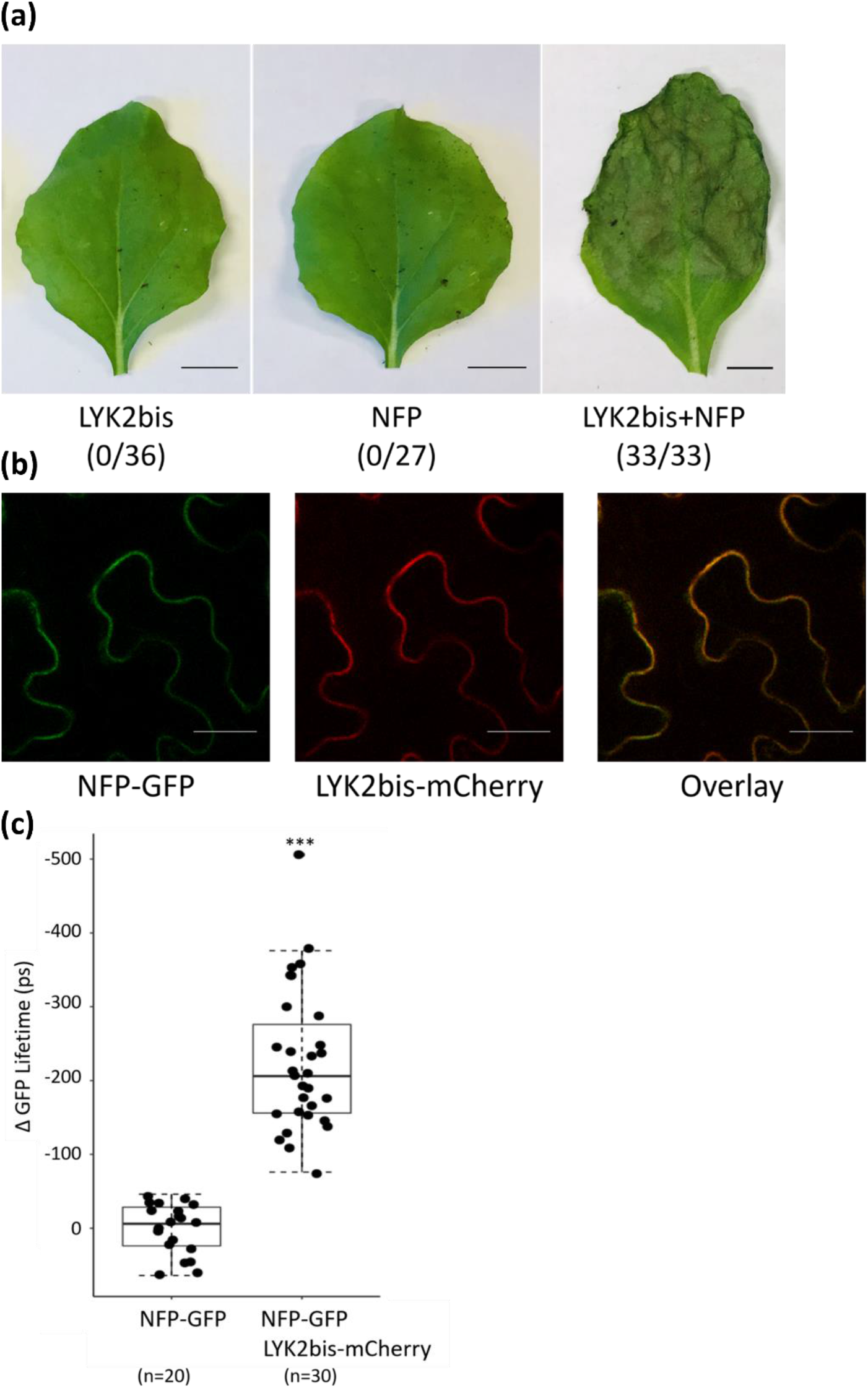
LYK2bis physically interacts with NFP and this interaction can lead to cell-death at the plasma membrane of *Nicotiana benthamiana* leaves. (a) Co-expression of LYK2bis and NFP induces cell-death in *N. benthamiana* leaves at 3 dpi. Numbers in brackets indicate the number of leaves exhibiting cell death/total number of infiltrated leaves. Bars, 1 cm. (b) Co-localisation of NFP and LYK2bis at the plasma membrane of *N. benthamiana* leaves. Leaf discs were observed by confocal microscopy at 2 dpi. Bars, 25 μm. (c) LYK2bis physically interacts with NFP at the plasma membrane of *N. benthamiana* leaves by FRET FLIM technology. For each measurement, the difference to average GFP decay times (Δ) of the GFP-NFP in the absence of the LYK2bis-mCherry is given in ps; n: number of samples measured for each condition. Students’ *t*-test was used for statistical analyses (***, *P* < 0.001).

At 2 dpi, the cell death response was much reduced and confocal microscopy revealed that the two proteins co-localise at the periphery of the cells (Fig. **8b**). As NFP was previously reported to localise at the plasma membrane of *N. benthamiana* leaves (Lefebvre *et al*., 2012), this evidence indicates that LYK2bis co-localises with NFP at the plasma membrane.

Similar leaf samples were used for FRET-FLIM analysis in which NFP-GFP was used as the donor and LYK2bis-mCherry was the acceptor. A highly significant decrease in lifetime of NFP-GFP was obtained when co-expressed with LYK2bis-mCherry (Fig. **8c**). This data not only supports the co-localization of LYK2bis and NFP at the plasma membrane but also provides strong evidence for the physical interaction between the two proteins.

### LYK2bis possesses an active kinase and can transphosphorylate the pseudo kinase of NFP

NFP has an inactive kinase domain (KD) which can be weakly trans-phosphorylated by the active kinase domain of LYK3 (Arrighi *et al*., 2006; Fliegmann *et al*., 2016). To test whether LYK2bis has an active kinase domain and can trans-phosphorylate NFP, we performed *in vitro* phosphorylation assays on the intracellular regions of the two proteins (which includes the KD), purified after expression in *E. coli*. We also tested the ability of LYK2bis-KD to trans-phosphorylate related proteins, LYR2-KD, LYR3-KD, LYR4-KD and LYK3-deadKD (Fliegmann *et al*., 2016), and the model kinase substrate myelin basic protein (MyBP). LYK2bis-KD had strong autophosphorylation activity and could trans-phosphorylate GST fusions of NFP-KD, LYR2-KD, LYR3-KD, LYR4-KD and LYK3-deadKD and MyBP in presence of [γ-^32^P] ATP, but not GST (Fig. **9**). These results show that LYK2bis has an active kinase domain and can trans-phosphorylate NFP, some other kinase domains and the model kinase substrate, MyBP.

**Fig. 9.**
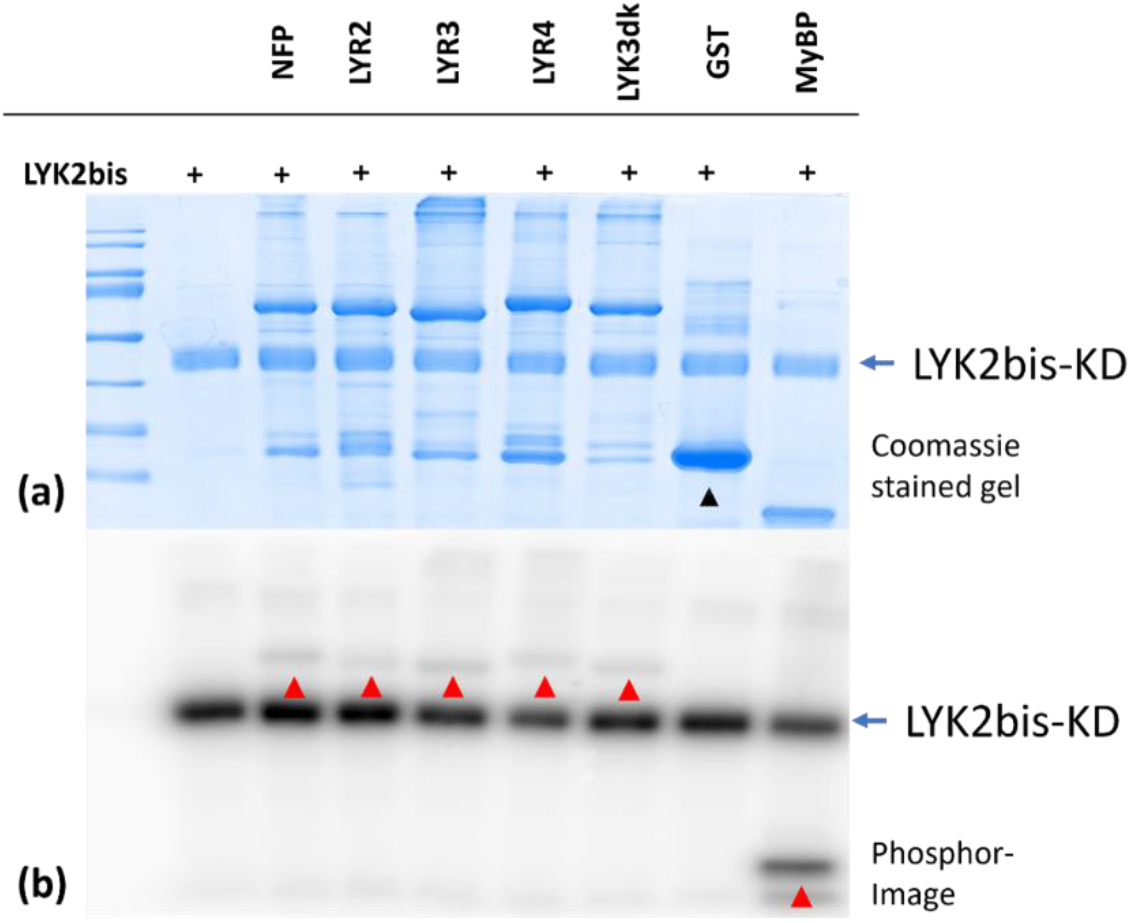
LYK2bis has an active kinase and can transphosphorylate NFP-KD *in vitro*. The kinase domains (KD) of LYK2bis and NFP were expressed and purified from *E. coli* as fusions with GST. GST-cleaved LYK2bis-KD was incubated individually or co-incubated with purified GST/NFP-KD (NFP), GST/LYR2-KD (LYR2), GST/LYR3-KD (LYR3), GST/LYR4-KD (LYR4), GST/LYK3-deadKD (LYK3dk), GST or Myelin Basic Protein (MyBP) in the presence of radioactive [γ-^32^P] ATP. Assays were analysed by SDS-PAGE, followed by coomassie staining (a) and phosphor imaging (b). Transphosphorylated proteins are marked by red arrowheads on the phosphor-image. The position of GST is marked on the coomassie gel (black arrowhead).

## Discussion

The symbiotic interactions between legumes and rhizobia are highly specific and require proper recognition of bacterial signals by plant receptors. NFs are key determinants of host specificity due to their species-specific chemical substitutions on both the reducing and non-reducing ends of the NF, which are recognized by receptors on roots of their compatible hosts. In this study, we demonstrate that two commonly used ecotypes of *M. truncatula*, A17 and R108, display a contrasting nodulation specificity with the *S. meliloti nodF/nodL* mutant; while A17 has a completely Nod^-^ phenotype with this strain, R108 is able to form infected nodules with both the *S. meliloti* WT and *nodF/nodL* strains (Fig. **1**). Using reverse genetics, we have identified *LYK2bis* as the genetic determinant underlying this extension of nodulation specificity. Complementation studies have clearly confirmed the essential role of *LYK2bis* in nodulation with the *nodF/nodL* strain of R108 (Fig. **3**). Moreover, this exceptional characteristic can be transferred to another genotype. As shown in Fig. **4**, the introduction of *LYK2bis* into A17 allows the plants to form nodules with the *nodF/nodL* mutant strain. These results provide conclusive evidence for the role of *LYK2bis* in nodulation with the *nodF/nodL* strain in R108.

### The evolution of LYK2bis

*LYK2bis* is predicted to encode a *LysM-RLK* and is located between *LYK2* and *LYK3* on chromosome 5 of the *M. truncatula* R108 genome (Fig. **2**). It is a chimeric gene with most of the LysM domains of *LYK2* and the rest, including the kinase domain, of *LYK3* (Fig. **S2**). Such a gene does not occur in other legumes such as *L. japonicus*, soybean and pea (De Mita *et al*., 2014; Sulima *et al*., 2017). In *M. truncatula*, although all the 23 genotypes available for BLAST screening in the Hapmap collection (https://medicagohapmap2.org/) contain sequences highly related to *LYK2* and *LYK3* (Table **S3**), only R108 contains *LYK2bis*. R108 is the most distant of the *M. truncatula* genomes (Zhou *et al*., 2017), and the only one representing *M. truncatula* ssp. *tricycla*. The most parsimonious explanation for the presence of *LYK2bis* in R108 is that it is a newly evolved gene formed from the pre-existing *LYK2* and *LYK3* genes in this genotype/subspecies, although we cannot exclude the possibility that it pre-existed in the most recent common ancestor of *M. truncatula* and has been lost at an early stage of the divergence of the two *M. truncatula* subspecies. In either case its phenotypic attributes suggest that it is a good example of recent adaptive gene duplication (Kondrashov, 2012).

### The specificity of nodulation directed by LYK2bis

Comparison of nodulation of the *lyk2bis* mutant with different *nod* gene mutants (Fig. **5**) suggest that it is the mutation in *nodL* (which is polar on *noeA, noeB*), and not in *nodF*, which is the major cause of lack of nodulation of the *lyk2bis* mutant by the *nodF/nodL* strain. Although the roles of *noeA* and *noeB* remain unknown, it is clear that NodL encodes an *O*-acetyl transferase and that the only difference detected in the NFs produced by this *nodL* mutant is the complete lack of *O*-acetylation of the non-reducing sugar (Ardourel et al., 1995). This evidence indicates a specific role of *LYK2bis* in nodulation with strains producing non-*O-*acetylated NFs whereas other gene(s) in R108 can enable nodulation with strains producing *O*-acetylated NFs, including ones without the specific C16 acyl chain, specified by the *nodFE* mutant. A17 shows only a slightly reduced nodulation with the *nodFE* mutant but a much reduced nodulation with a *nodL* mutant, and this has been attributed to the *LYK3* gene (Smit *et al*., 2007). LYK3-A17 may thus show a preference for recognizing *O*-acetylated NFs but may tolerate C18 ones. It is relevant to note that in pea, nodulation with C18:1 NFs produced by a *Rhizobium leguminosarum nodE* mutant is associated with certain haplotypes of *PsSYM37*, which is orthologous and functionally equivalent to *LYK3/NFR1* (Li *et al*., 2011).

In pea, an interesting phenotype linked to *O*-acetylation of the reducing sugar of NFs has been identified in cv Afghanistan: nodulation by *R. leguminosarum* requires this particular NF structural modification, which is produced by strains containing the *nodX* gene (Firmin *et al*., 1993). On the plant side, *PsSym2* was identified as the determinant of this strain selectivity, and incidentally, synteny with closely related *M. truncatula* led to the identification of the *LYK* cluster, containing genes *LYK1* -*LYK7* (Limpens *et al*., 2003). Recent studies have identified three *LysM-RLKs* in the corresponding pea *LYK* cluster: *PsSYM37, PsK1* and *PsLYKX*, all of which play roles in nodulation (Zhukov *et al*., 2008; Li *et al*., 2011; Kirienko *et al*., 2019; Sulima *et al*., 2017). By analysing different pea cultivars, it has been shown that nodulation with *nodX*^+^ strains is correlated with haplotypes of the *PsLYKX* gene, but not *PsSYM37* or *PsK1*, suggesting that *PsLYKX* corresponds to *Sym2* (Sulima *et al*., 2017, 2019).

### The importance of LYK2bis structure for NF recognition

Recently, Bozsoki *et al*. (2020) have published the crystal structure of the extracellular domain of LYK3-A17. By using a domain swapping and point mutation approach, the authors have identified the importance of two regions in the LysM1 domain of LYK3-A17 for the specificity of NF signalling. As shown in Fig. **S1**, the extracellular domain of LYK2bis shows strong divergence to LYK3-A17, especially in LysM1 (Table **S2**, Table **S5**). In particular, the two regions II and IV of LysM1 that are essential in LYK3 for discrimination of NF and CO ligands and for specific nodulation are very poorly conserved in LYK2bis. These regions could therefore be involved in the discrimination of *nodF/nodL* and WT NFs. In pea, nodulation with *nodX*^+^ strains is associated with *LYKX* haplotypes containing specific amino acids in regions II and IV of LysM1, as described above (Sulima *et al*., 2019; Solovev *et al*., 2021). Together, this evidence indicates the importance of specific regions in the LysM1 of these highly-related LYK cluster proteins in the recognition of specific NF decorations.

### Mechanism of action of LYK2bis

MtNFP, and its orthologs in *L. japonicus* (NFR1), *P. sativum* (SYM10) and other legumes, is the key LysM-RLK involved in NF perception and signalling (Krönauer & Radutoiu, 2021). Many studies have shown the involvement of this protein and one or two LYK cluster proteins in perception of NFs and signalling for nodulation (Radutoiu *et al*., 2003; Kirienko *et al*., 2019). In this study, we observed that the co-expression of LYK2bis and NFP in *N. benthamiana* leaves leads to defence-like responses, which were also observed in co-expression of LYK3-A17 and NFP (Pietraszewska-Bogiel *et al*., 2013), indicating a similar mechanism of action between these two protein pairs. In addition, we have shown that LYK2bis physically interacts with NFP in *N. benthamiana* leaves suggesting the formation of a heteromer *in planta*, which is possibly responsible for recognition of non-*O*-acetylated NFs. This hypothesis is supported by the work on the interaction between PsLYKX and PsSYM10 receptor complex with NFs (Solovev *et al*., 2021). In this study, using molecular modelling and ligand docking, the authors have shown the potential for stable heterodimers between PsLYKX and PsSYM10, which may interact with NFs at the heterodimer interface. Indeed in LYK3, region III of LysM1 is envisaged to interact with NFP (Bozsoki *et al*., 2020), and is quite highly conserved between LYK2bis and LYK3 (7/12 residues – Fig. **S2**). Moreover, the LYK2bis intracellular domain is almost identical to that of LYK3-A17 (three amino acids in difference) and both possess an active kinase, which *in vitro* can trans-phosphorylate NFP-KD, albeit non-specifically (Fig. **8** and Fliegmann *et al*., 2016). This similarity suggests a mechanism by which LYK2bis could integrate into the LYK3 signal transduction pathway. This may be through substitution for LYK3, rather than interaction with this protein, as we have shown that the LYK2bis-dependent gain of A17 nodulation by the *nodF/nodL* strain can occur in a *lyk3* mutant.

Finally, our work has shown that *LYK2bis* is very important for nodulation of R108 by many natural strains of *S. meliloti* and *S. medicae*. However, some strains of each species do not require this gene for R108 nodulation (Table **S4** and Fig. **6**). This difference in *LYK2bis* dependence could be explained by the strains differing in the proportion of non-*O*-acetylated/*O*-acetylated NFs that they produce: the dependent strains producing a greater proportion than the *LYK2bis*-independent strains. Analysis of the *nodL* gene from the available sequences of these strains did not reveal any differences that correlated with the two classes (B. Gourion, personal communication). However, recent preliminary analysis of the NFs from *S. medicae* WSM419 supports this hypothesis as *O*-acetylated NFs were not detected among the most abundant detected NFs (V. Puech*-*Pagès, F. Maillet, B. Gourion, P. Ratet, personal communication), whereas the *LYK2bis* independent strain *S. meliloti* 2011 produces a majority of *O*-acetylated NFs (Roche et al., 1991). R108 was collected from Israel (Garmier *et al*., 2017) but no correlation was observed between *LYK2bis*-dependence and strains from the Middle East region, suggesting that *LYK2bis* has not evolved to adapt to a particular strain, specific to this geographical region. Genomic studies on *S. meliloti* and *S. medicae* suggest a complex history of horizontal gene transfer between these two species, which is restricted almost exclusively to plasmid genes, including the *nod* genes (Epstein *et al*., 2012). Thus, the ability to produce non-*O*-acetylated NFs may be widespread but patchy in *Sinorhizobium* and the evolution of *LYK2bis* may thus present an adaptive advantage to allow nodulation by a greater variety of strains.

Based on data obtained in this study, we thus propose a model for the nodulation of R108 in which rhizobia may produce different proportions of *O*-acetylated/non-*O*-acetylated NFs. LYK2bis forms a receptor complex with NFP that perceives non-*O*-acetylated NFs and activates the kinase of LYK2bis, which then trans-phosphorylates NFP leading to downstream signalling and finally to nodule formation (Fig. **10**). Other *LYK* genes such as *LYK3*, possibly redundant with *LYK2bis*, may be involved in the perception of O-acetylated NFs.

**Fig. 10.**
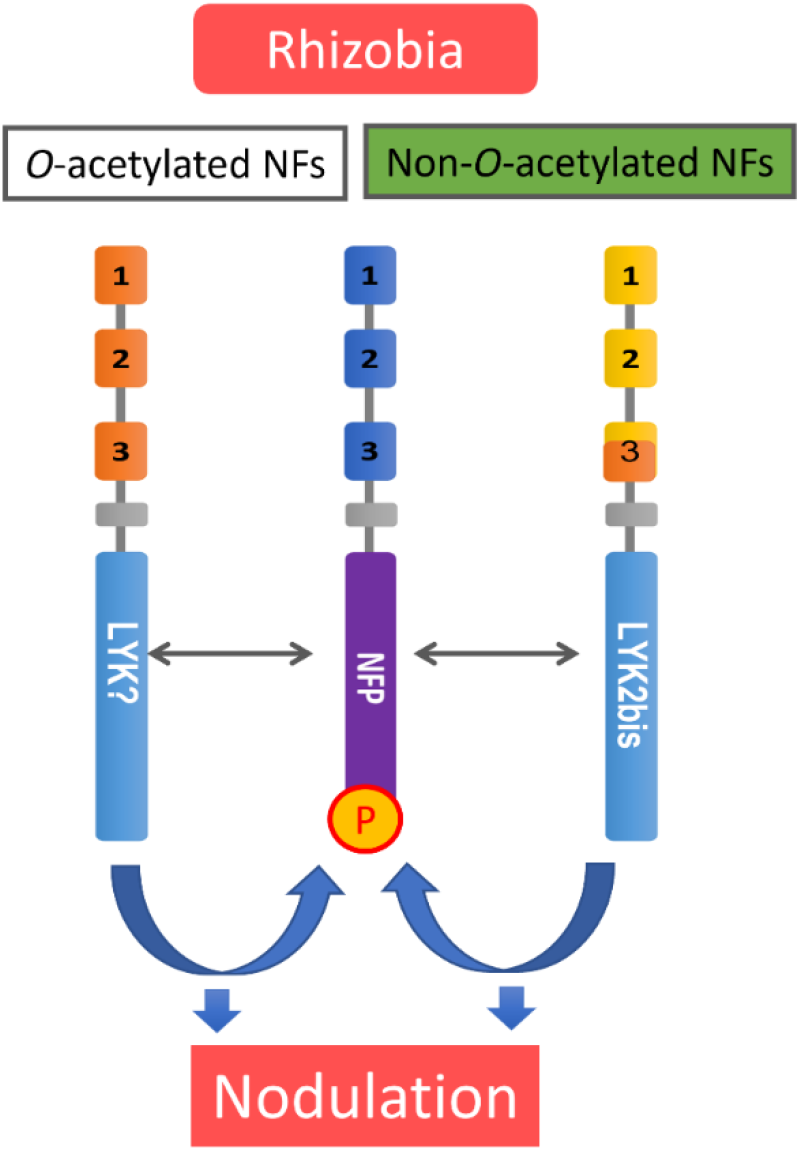
Model of NF recognition and nodulation in *Medicago truncatula* R108. Different rhizobia may secrete different proportions of *O*-acetylated and non-*O*-acetylated NFs. The nodulation of R108 with strains producing non-*O*-acetylated NFs is dependent on LYK2bis and possibly involves NFP while the *O*-acetylated NFs may involve other LYK(s). The active LYK kinases can transphosphorylate (P) NFP, which may be important for signal transduction.

In conclusion, we have identified a newly-evolved gene in R108, *LYK2bis*, which is required for efficient nodulation by many but not all *Sinorhizobium meliloti and medicae* strains and is responsible for extending the nodulation specificity of R108 to include a *nodF/nodL* mutant. *LYK2bis* appears to have a specific role in recognising NFs which are non-*O*-acetylated on the terminal non-reducing sugar. *LYK2bis* is a chimera formed from the *LYK2* and *LYK3* genes and is located between them in the *LYK* cluster on chromosome 5. Studies on the diversity of this locus and the expansion of the *tricycla* subspecies, coupled with analysis of rhizobial symbionts, would establish whether the evolution of this gene in *M. truncatula* has led to host-range expansion.

## Acknowledgements

We thank Clare Gough, Sandra Bensmihen, Benoit Lefebvre, Benjamin Gourion (LIPME) and Pierre-Marc Delaux and Maxime Bonhomme (Laboratoire de Recherche en Sciences Végétales, Toulouse) for discussions and for their comments on the manuscript. We are very grateful to Michael Sadowsky, Peter Tiffin (University of Minnesota, USA), Katy Heath (University of Illinois, USA) and Gilles Bena (UMR Interactions Plantes-Microorganismes-Environnement, Montpellier, France) for discussions on Rhizobium and sending us the strains. We thank Virginie Puech*-*Pagès (LRSV), Fabienne Maillet (LIPME), Benjamin Gourion (LIPME) and Pascal Ratet (IPS2) for allowing us to cite their Nod factor analysis and Benjamin Gourion (LIPME) for analysis of the *nodL* gene from selected *Sinorhizobium* genotypes. We thank Fabienne Maillet and Virginie Gasciolli (LIPME) for technical advice. For the *Tnt1* mutants, the *Medicago truncatula* plants utilized in this research project, which are jointly owned by the Centre National de la Recherche Scientifique, were obtained from Noble Research Institute, LLC and were created through research funded, in part, by a grant from National Science Foundation, NSF-0703285. Funding for part of this work was gratefully received from the Fédération de Recherche Agrobiosciences, Interactions et Biodiversité (FR AIB – project CHAIN, coordinators J. Cullimore and C. Jacquet) and the Agence Nationale de la Recherche (ANR – project DUALITY, ANR-20-CE20-0017-01, coordinator C. Gough). This study is set within the framework of the “Laboratoires d’Excellences (LABEX)” TULIP (ANR 10 LABX 41) and of the “École Universitaire de Recherche (EUR)” TULIP GS (ANR 18 EURE 0019). TBL gratefully acknowledges receipt of a Bourse d’Excellence from the Ambassade de France au Vietnam, to fund her PhD in France.

## Author contribution

TBL, NP and JC designed and performed experiments, interpreted the data and wrote the manuscript. AO contributed to experiments on natural rhizobium strains and performed the mycorrhization test. CP performed and analysed the FRET-FLIM experiments.

## Supporting Information

Additional supporting information may be found in the online version of this article.

**Fig. S1** *Medicago truncatula* R108 is able to form fixing nodules with both *Sinorhizobium meliloti* 2011 WT and its *nodF/nodL* mutant.

**Fig. S2** Amino acid alignment of LYK3-A17, LYK3-R108, LYK2bis, LYK2-A17 and LYK2-R108

**Table S1** *Sinorhizobium meliloti* 2011 strains used in this study

**Table S2** Percentage of identity/similarity of LYK2, LYK3 in A17 and LYK2bis and LYK3 in R108

**Table S3** Presence of *LYK2bis* in *Medicago truncatula* genomes

**Table S4** Characteristics of natural *Sinorhizobium* strains and their ability to nodulate R108 and the *lyk2bis-1R* mutant.

**Table S5** Number of identical amino acids/total amino acids in each LysM between LYK2bis and LYK3-A17 and LYK2-A17

