## Supporting Information for "A newly-evolved chimeric lysin motif receptor-like kinase in *Medicago truncatula* spp. *tricycla* R108 extends its Rhizobia symbiotic partnership"

Article acceptance date: Click here to enter a date.

The following Supporting Information is available for this article:

**Fig. S1** *Medicago truncatula* R108 is able to form fixing nodules with both *Sinorhizobium meliloti* 2011 WT and its *nodF/nodL* mutant.

**Fig. S2** Amino acid alignment of LYK3-A17, LYK3-R108, LYK2bis, LYK2-A17 and LYK2-R108

**Table S1** *Sinorhizobium meliloti* 2011 strains used in this study

**Table S2** Percentage of identity/similarity of LYK2, LYK3 in A17 and LYK2bis and LYK3 in R108

**Table S3** Presence of *LYK2bis* in *Medicago truncatula* genomes

**Table S4** Characteristics of natural *Sinorhizobium* strains and their ability to nodulate R108 and the *lyk2bis-1R* mutant

**Table S5** Number of identical amino acids/total amino acids in each LysM between LYK2bis and LYK3-A17 and LYK2-A17

**Fig. S1** *Medicago truncatula* R108 is able to form fixing nodules with both *Sinorhizobium meliloti* 2011 WT and its *nodF/nodL* mutant. The nitrogenase activity of R108 were measured in 16 plants/inoculation at 28dpi. Statistical analyses were performed using Student's *t*-test (ns, not significant). White shading is for plants inoculated with the WT strain; green shading is for plants incoulated with the *nodF/nodL* strain.

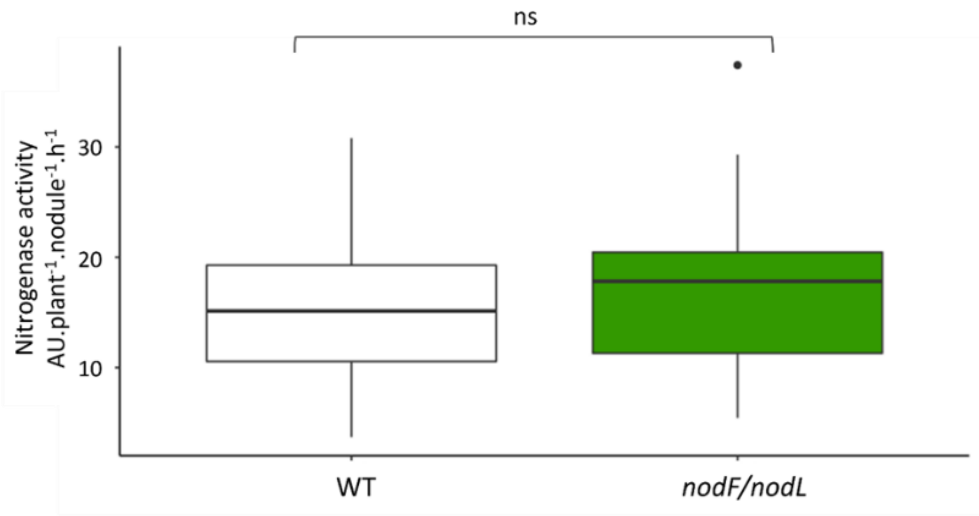

**Fig. S2** Amino acid alignment of LYK3-A17, LYK3-R108, LYK2bis, LYK2-A17 and LYK2-R108. The three LysM domains are indicated by green boxes following the crystal structure of LYK3-A17 extracellular domain (Bozsoki *et al.*, 2020). The three regions II, III and IV that are important for NF recognition in the LysM1 of LYK3-A17 are highlighted in yellow, as is the corresponding region of LYK2bis. Red arrows indicate the conserved CXC motifs separating LysM domains. The transmembrane (TM), and kinase domain (KD) are marked in blue and purple, respectively.

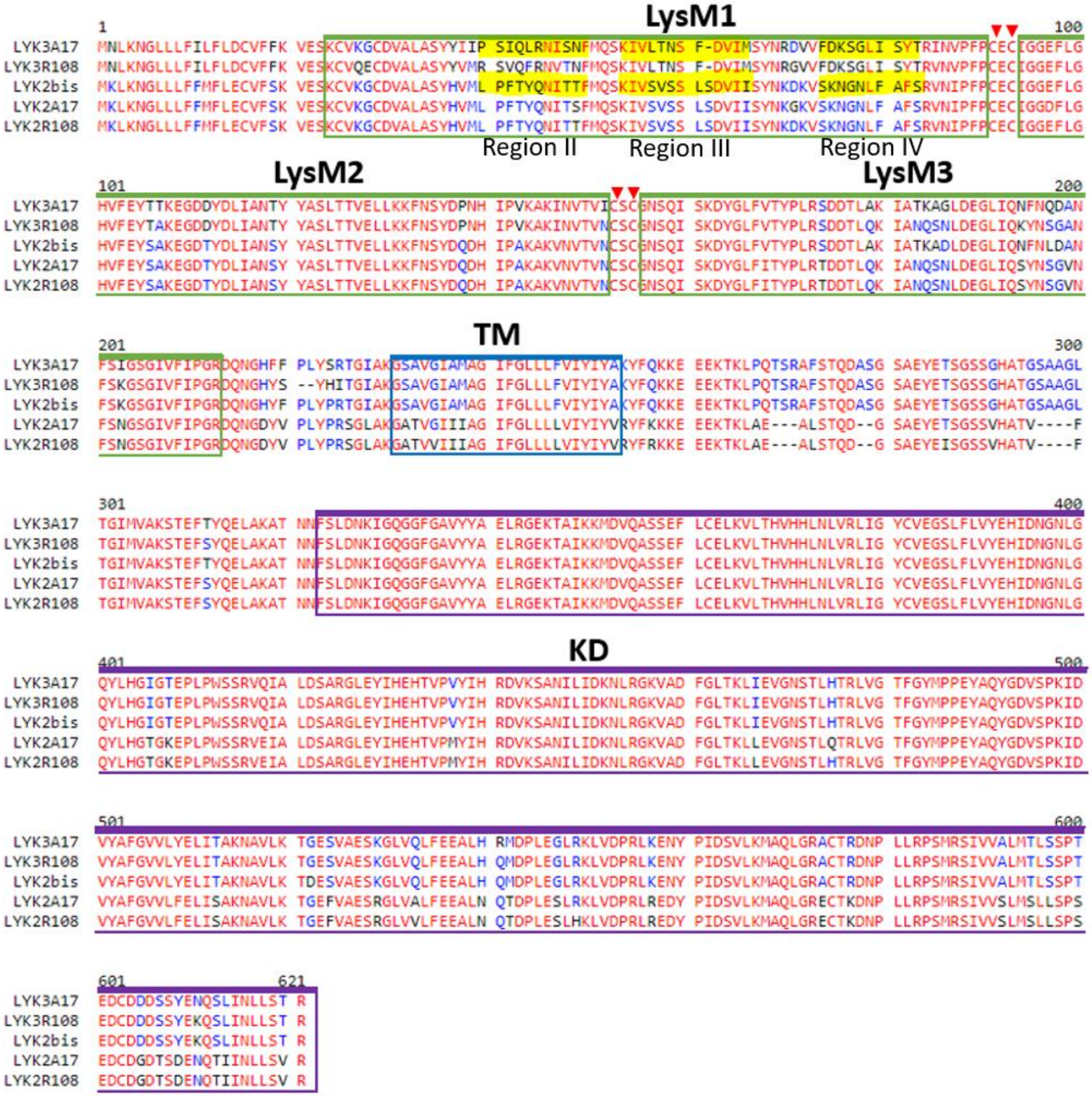

**Table S1** *Sinorhizobium meliloti* mutant strains used in this study

| Designation | | Relevant characteristics | Reference |
|---|---|---|---|
| *S. meliloti* | | | |
| | GMI6526 | 2011(pXLGD4), Nod+Fix+ on *M. truncatula* | Ardourel *et al*., 1994 |
| | GMI6630 | 2011Δ(nodF) nodL::Tn5 (pXLGD4) | Ardourel *et al*., 1994 |
| | GMI6528 | 2011Δ(nodF) Δ(nodE) (pXLGD4) | Ardourel *et al*., 1994 |
| | GMI6563 | 2011 nodL::Tn5 (pXLGD4) | Ardourel *et al*., 1994 |

**Table S2** Percentage identity/similarity of LYK2, LYK3 in A17 and LYK2, LYK2bis and LYK3 in R108. Data were calculated using EMBOSS Needle for the whole proteins and for the predicted extracellular (ECD) and intracellular (ICD) domains. The "heat map" colouring indicates the degree of identity and similarity.

| | whole protein | | ECD | | ICD | |
|---|---|---|---|---|---|---|
| | LYK2-A17 | LYK-3A17 | LYK2-A17 | LYK3-A17 | LYK2-A17 | LYK-3A17 |
| **Identity** | | | | | | |
| **LYK2-R108** | 98.5 | 81.0 | 98.7 | 72.1 | 98.6 | 87.1 |
| **LYK2bis** | 88.1 | 91.6 | 90.6 | 78.5 | 87.1 | 99.2 |
| **LYK3R108** | 82.8 | 94.8 | 75.5 | 87.5 | 87.7 | 99.2 |
| **Similarity** | | | | | | |
| **LYK2-R108** | 99.0 | 89.2 | 99.6 | 84.5 | 98.9 | 92.6 |
| **LYK2bis** | 93.4 | 95.0 | 95.7 | 87.1 | 92.3 | 99.5 |
| **LYK3-R108** | 90.0 | 96.9 | 86.3 | 92.2 | 92.6 | 99.7 |

**Table S3** Presence of *LYK2bis* in *Medicago truncatula* genomes. The 1st exon of *LYK2, LYK3* and *LYK2bis* from R108 were used to blast on all genomes available in the Hapmap2 database and the first hit for each query is shown. The percentages of identity (%ID) between hits found with the *LYK3-R108* query and the *LYK2bis* sequence were calculated by EMBOSS Needle.

| ***Medicago truncatula*** **genomes** | **First hit with LYK2R108 and LYK2bis queries** | **Positions** | | **%ID to LYK2R108** | **%ID to LYK2bis** | **First hit with LYK3R108 query** | **Positions** | | **%ID to LYK3R108** | **%ID to LYK2bis** |
|---|---|---|---|---|---|---|---|---|---|---|
| | | **Sstart** | **Send** | | | | **Sstart** | **Send** | | |
| HM002 | medtr.HM002.gnm1.scaffold_66 | 1147008 | 1147647 | 99.1 | 96.3 | medtr.HM002.gnm1.scaffold_66 | 1088106 | 1088742 | 93.6 | 82.9 |
| HM004 | medtr.HM004.gnm1.scaffold_49 | 757127 | 756488 | 99.1 | 96.6 | medtr.HM004.gnm1.scaffold_49 | 791170 | 790534 | 97.2 | 81.0 |
| HM005 | medtr.HM005.gnm1.scaffold_247 | 74669 | 74030 | 98.0 | 95.2 | medtr.HM005.gnm1.scaffold_247 | 138099 | 137463 | 94.2 | 82.0 |
| HM006 | medtr.HM006.gnm1.scaffold_251 | 319609 | 320248 | 98.6 | 95.8 | medtr.HM006.gnm1.scaffold_251 | 286050 | 286683 | 94.6 | 82.6 |
| HM010 | medtr.HM010.gnm1.scaffold_29 | 80819 | 80180 | 99.1 | 96.3 | medtr.HM010.gnm1.scaffold_29 | 130689 | 130053 | 94.5 | 82.6 |
| HM017 | medtr.HM017.gnm1.scaffold_27 | 30550 | 31083 | 99.1 | 99.1 | medtr.HM017.gnm1.scaffold_128 | 725038 | 725674 | 93.1 | 83.1 |
| HM018 | medtr.HM018.gnm1.scaffold_309 | 71485 | 70843 | 97.2 | 94.7 | medtr.HM018.gnm1.scaffold_309 | 110156 | 109520 | 93.9 | 83.4 |
| HM020 | medtr.HM020.gnm1.scaffold_57 | 487808 | 488447 | 97.5 | 95.0 | medtr.HM020.gnm1.scaffold_57 | 431300 | 431933 | 97.3 | 80.7 |
| HM022 | medtr.HM022.gnm1.scaffold_48 | 230635 | 231274 | 98.0 | 95.2 | medtr.HM022.gnm1.scaffold_48 | 197372 | 198008 | 93.7 | 83.4 |
| HM023 | medtr.HM023.gnm1.scaffold_313 | 72441 | 71799 | 97.2 | 94.7 | medtr.HM023.gnm1.scaffold_313 | 110985 | 110349 | 93.9 | 83.4 |
| HM026 | medtr.HM026.gnm1.scaffold_781 | 53781 | 53142 | 98.8 | 96.9 | medtr.HM026.gnm1.scaffold_781 | 95385 | 95010 | 96.3 | 46.3 |
| HM034 | medtr.HM034.gnm1.scaffold_30 | 30716 | 30077 | 99.1 | 96.3 | medtr.HM034.gnm1.scaffold_30 | 74077 | 73441 | 93.6 | 83.4 |
| HM050 | medtr.HM050.gnm1.scaffold_33 | 235744 | 235105 | 97.0 | 94.5 | medtr.HM050.gnm1.scaffold_33 | 290656 | 290020 | 93.9 | 83.1 |
| HM056 | medtr.HM056.gnm1.scaffold_503 | 143651 | 144290 | 99.1 | 96.3 | medtr.HM056.gnm1.scaffold_503 | 106282 | 106915 | 94.3 | 82.5 |
| HM058 | medtr.HM058.gnm1.scaffold_687 | 107608 | 106966 | 96.7 | 94.3 | medtr.HM058.gnm1.scaffold_687 | 120748 | 120112 | 95.9 | 80.9 |
| HM060 | medtr.HM060.gnm1.scaffold_250 | 27295 | 26656 | 99.4 | 96.6 | medtr.HM060.gnm1.scaffold_250 | 60980 | 60344 | 97.2 | 81.0 |
| HM095 | medtr.HM095.gnm1.scaffold_274 | 80513 | 81152 | 99.1 | 96.3 | medtr.HM095.gnm1.scaffold_274 | 33305 | 33941 | 93.6 | 83.4 |
| HM125 | medtr.HM125.gnm1.scaffold_412 | 44341 | 44983 | 96.7 | 94.3 | medtr.HM125.gnm1.scaffold_412 | 31149 | 31785 | 95.9 | 80.9 |
| HM129 | medtr.HM129.gnm1.scaffold_250 | 82289 | 81650 | 99.2 | 96.4 | medtr.HM129.gnm1.scaffold_250 | 95260 | 94624 | 95.9 | 80.9 |
| HM185 | medtr.HM185.gnm1.scaffold_10 | 363827 | 364466 | 99.4 | 96.6 | medtr.HM185.gnm1.scaffold_10 | 320702 | 321338 | 95.9 | 80.9 |
| HM324 | medtr.HM324.gnm1.scaffold_870 | 39565 | 40204 | 99.2 | 96.4 | medtr.HM324.gnm1.scaffold_433 | 227646 | 228282 | 93.9 | 83.4 |
| HM340-R108 | medtr.R108_HM340.gnm1.scf013 | 1134685 | 1134046 | 96.9 | 100.0 | medtr.R108_HM340.gnm1.scf013 | 1153922 | 1153286 | 100.0 | 83.9 |
| HM340-R108 | medtr.R108_HM340.gnm1.scf013 | 1040902 | 1040263 | 100.0 | 96.9 | | | | | |
| HM341-A17 | medtr.A17_HM341.gnm4.chr5 | 37308676 | 37309315 | 98.8 | 96.3 | medtr.A17_HM341.gnm4.chr5 | 37236497 | 37237133 | 94.5 | 82.6 |

**Table S4** Characteristics of natural *Sinorhizobium* strains and their ability to nodulate R108 and the *lyk2bis-1R* mutant. Four to five plants per genotype were grown in tubes on agar slants and inoculated with 10,000 bacteria/plant of the corresponding Sinorhizobium strain. Nodulation was analysed at 31dpi.

| **Sinorhizobium strains** | | | | | **Nodulation phenotyping** | | | |
|---|---|---|---|---|---|---|---|---|
| **Species** | **Strain** | **Origin** | **Plant origin** | **Reference** | **Genotype** | **N° of nodules** | **Difference R108/ *lyk2bis*** | **% pink nodules** |
| *S. meliloti* | 2011 | | | LIPME | R108 | 11.7 | | 82 |
| | | | | | *lyk2bis* | 7.7 | | 93 |
| *S. meliloti* | 2011 *nodF/nodL* | | | LIPME | R108 | 11.8 | ** | 60 |
| | | | | | *lyk2bis* | 0 | | |
| *S. meliloti* | Rm41 | Hungary | *M. sativa* | Sugawara *et al.* , 2013 | R108 | 14 | * | 7 |
| | | | | | *lyk2bis* | 0 | | |
| *S. medicae* | WSM419 | Italy | *M. murex* | Reeve *et al.* , 2010 | R108 | 12.3 | * | 100 |
| | | | | | *lyk2bis* | 0 | | |
| *S. meliloti* | M162 | Syria | *M. truncatula* | Sugawara *et al.* , 2013 | R108 | 16 | * | 94 |
| | | | | | *lyk2bis* | 4.5 | | 61 |
| *S. meliloti* | M270 | Jordan | *M. truncatula* | Sugawara *et al.* , 2013 | R108 | 21.3 | * | 0 |
| | | | | | *lyk2bis* | 7.3 | | 0 |
| *S. meliloti* | M156 | Syria | *M. rigidula* | Sugawara *et al.* , 2013 | R108 | 18.5 | ** | |
| | | | | | *lyk2bis* | 2.8 | | |
| *S. meliloti* | M195 | Turkey | *M. rigidula* | Sugawara *et al.* , 2013 | R108 | 14 | NSD | 100 |
| | | | | | *lyk2bis* | 8.8 | | 74 |
| *S. meliloti* | M10 | Syria | *M. blancheana* | Sugawara *et al.* , 2013 | R108 | 21.3 | * | 0 |
| | | | | | *lyk2bis* | 0.5 | | 0 |
| *S. meliloti* | T027 | Tunisia | *M. truncatula* | Sugawara *et al.* , 2013 | R108 | 14.3 | NSD | 93 |
| | | | | | *lyk2bis* | 10.5 | | 100 |
| *S. medicae* | M22 | Syria | *M. polymorpha* | Sugawara *et al.* , 2013 | R108 | 5.3 | * | 69 |
| | | | | | *lyk2bis* | 1.3 | | 100 |
| *S. medicae* | M58 | Jordan | *M. rotata* | Sugawara *et al.* , 2013 | R108 | 16.3 | NSD | 100 |
| | | | | | *lyk2bis* | 12.8 | | 78 |
| *S. medicae* | M102 | Syria | *M. truncatula* | Sugawara *et al.* , 2013 | R108 | 9.5 | ** | 100 |
| | | | | | *lyk2bis* | 1.5 | | 0 |
| *S. saheli* | USDA 4893 | Senegal | *Sesbania cannabina* | Sugawara *et al.* , 2013 | R108 | 12 | * | 79 |
| | | | | | *lyk2bis* | 0 | | |
| *S. terangae* | USDA 4894 | Senegal | *Acacia laeta* | Sugawara *et al.* , 2013 | R108 | 0 | | |
| | | | | | *lyk2bis* | 0 | | |
| *S. meliloti* | STM2775 | France | *M. littoralis* | Bailly *et al.* , 2006 | R108 | 12.2 | * | 92 |
| | | | | | *lyk2bis* | 5.6 | | 86 |
| *S. meliloti* | STM1661 | France | *M. littoralis* | Bailly *et al.* , 2006 | R108 | 11.3 | * | 80 |
| | | | | | *lyk2bis* | 0 | | |
| *S. meliloti* | STM2772 | France | *M. rigiduloides* | Bailly *et al.* , 2006 | R108 | 12.8 | NSD | 100 |
| | | | | | *lyk2bis* | 7 | | 80 |
| *S. meliloti* | STM1643 | France | *M. truncatula* | Bailly *et al.* , 2006 | R108 | 15.3 | NSD | 82 |
| | | | | | *lyk2bis* | 4.3 | | 100 |
| *S. meliloti* | STM2748 | France | *M. truncatula* | Bailly *et al.* , 2006 | R108 | 19.2 | * | 11 |
| | | | | | *lyk2bis* | 7.4 | | 0 |
| *S. medicae* | STM 2745 | France | *M. truncatula* | Bailly *et al.* , 2006 | R108 | 16.8 | NSD | 75 |
| | | | | | *lyk2bis* | 4.8 | | 100 |
| *S. medicae* | STM1647 | France | *M. ciliaris* | Bailly *et al.* , 2006 | R108 | 6.4 | *** | 91 |
| | | | | | *lyk2bis* | 1.2 | | 83 |
| *S. medicae* | STM1654 | France | *M. polymorpha* | Bailly *et al.* , 2006 | R108 | 14.4 | * | 54 |
| | | | | | *lyk2bis* | 0.7 | | 0 |
| *S. medicae* | STM2744 | France | *M. truncatula* | Bailly *et al.* , 2006 | R108 | 3.8 | NSD | 100 |
| | | | | | *lyk2bis* | 0 | | |
| Mock | | | | | R108 | 0 | | |
| | | | | | *lyk2bis* | 0 | | |

NSD = No significant difference ($P > 0.05$). ***, $P < 0.001$; **, $P < 0.01$; *, $P < 0.05$

**Table S5** Number of identical amino acids/total amino acids in each LysM between LYK2bis and LYK3-A17 and LYK2-A17

| LYK2bis | Identities with LYK3A17 | Identities with LYK2A17 |
|---|---|---|
| **LysM1** | 40/67 | 65/67 |
| **LysM2** | 52/59 | 59/59 |
| **LysM3** | 55/58 | 51/58 |
